# Single-Trial Decoding of Scalp EEG Under Natural Conditions

**DOI:** 10.1101/481630

**Authors:** Greta Tuckute, Sofie Therese Hansen, Nicolai Pedersen, Dea Steenstrup, Lars Kai Hansen

## Abstract

There is significant current interest in decoding mental states from electro-encephalography (EEG) recordings. EEG signals are subject-specific, sensitive to disturbances, and have a low signal-to-noise ratio, which has been mitigated by the use of laboratory-grade EEG acquisition equipment under highly controlled conditions. In the present study, we investigate single-trial decoding of natural, complex stimuli based on scalp EEG acquired with a portable, 32 dry-electrode sensor system in a typical office setting. We probe generalizability by a leave-one-subject-out cross-validation approach. We demonstrate that Support Vector Machine (SVM) classifiers trained on a relatively small set of de-noised (averaged) pseudo-trials perform on par with classifiers trained on a large set of noisy single-trial samples. For visualization of EEG signatures exploited by SVM classifiers, we propose a novel method for computing sensitivity maps of EEG-based SVM classifiers. Moreover, we apply the NPAIRS resampling framework for estimation of map uncertainty and show that effect sizes of sensitivity maps for classifiers trained on small samples of de-noised data and large samples of noisy data are similar. Finally, we demonstrate that the average pseudo-trial classifier can successfully predict the class of single trials from withheld subjects, which allows for fast classifier training, parameter optimization and unbiased performance evaluation in machine learning approaches for brain decoding.

## 1. INTRODUCTION

Decoding of brain activity aims to predict the perceptual and semantic content of neural processing based on activity measured in one or more brain imaging modalities, such as electro-encephalography (EEG), magneto-encephalography (MEG), and functional magnetic resonance imaging (fMRI). Decoding studies based on fMRI have matured significantly during the last 15 years (see e.g. [Haynes and Rees, 2006, Gerlach, 2007] for review), and human brain activity has been successfully decoded from natural images and movies [Kay et al., 2008, Prenger et al., 2009, Nishimoto et al., 2011, Huth et al., 2012, Huth et al., 2016, Güçlü and van Gerven, 2017].

In case of decoding of scalp EEG, the research area is still progressing, and relatively few studies document detection of brain states in regards to semantic categories (often discrimination between two high-level categories) [Simanova et al., 2010, Murphy et al., 2011, Wang et al., 2012, Taghizadeh-Sarabi et al., 2014, Stewart et al., 2014, Kaneshiro et al., 2015, Zafar et al., 2017]. EEG-based decoding of human brain activity has significant potential due to excellent time resolution and the possibility of real-life acquisition, however, the signal is extremely diverse, subject-specific, sensitive to disturbances, and has a low signal-to-noise ratio, hence, posing a major challenge for both signal processing and machine learning [Nicolas-Alonso and Gomez-Gil, 2012].

Due to before-mentioned challenges, previous studies have been performed in controlled laboratory settings with high-grade EEG acquisition equipment [Simanova et al., 2010, Murphy et al., 2011, Wang et al., 2012, Stewart et al., 2014, Kaneshiro et al., 2015, Zafar et al., 2017]. Visual stimuli paradigms can often not be described as naturalistic, due to 1) repeated presentation of identical experimental trials, and 2) iconic views of objects and lack of complexity of semantic context [Simanova et al., 2010, Murphy et al., 2011, Wang et al., 2012, Taghizadeh-Sarabi et al., 2014, Stewart et al., 2014, Kaneshiro et al., 2015]. Generalizability of decoding classifier models to novel participants is rare, due to subject-specific modelling approaches [Simanova et al., 2010, Wang et al., 2012, Stewart et al., 2014, Taghizadeh-Sarabi et al., 2014, Kaneshiro et al., 2015, Zafar et al., 2017]. Moreover, a number of participants are occasionally excluded from analysis due to artifacts and low classification accuracy [Taghizadeh-Sarabi et al., 2014, Zafar et al., 2017].

The motivation for the present study is to overcome the highlighted limitations in EEG-based decoding. The current experimental paradigm and decoding work is centered around 1) ecological validity and portability, and 2) generalizability. Therefore, we acquired scalp EEG signals in a typical office setting using a portable, user-friendly, wireless EEG Enobio system with 32 dry electrodes. Experimental image stimuli consisted of non-iconic views of objects embedded in complex everyday scenes (Figure 1A) of 23 different semantic categories from an open image database [Lin et al., 2014]. All images presented were unique and not repeated for the same subject throughout the experiment (Figure 1B), akin to how visual stimuli are experienced in real life. We created classifiers based on single-trial responses as well as generalized category representations by averaging responses of images from the same semantic category.

We acquired data from 15 healthy participants (5 female). We are interested in exploring the limitations of inter-subject generalization, i.e., population models, hence no participants are excluded from analysis. Decoding ability is evaluated in an inter-subject design, i.e., in a leave-one-subject-out approach (as opposed to within-subject classification) to probe generalizability across participants [Kjems et al., 2002].

The work in the present study is focused on the binary classification problem between two classes: Brain processing of animate and inanimate image stimuli. Kernel methods, e.g., support vector machines (SVM) are frequently applied for learning of statistical relations between patterns of brain activation and experimental conditions. In classification of EEG data, SVMs have shown good performance in many contexts [Murphy et al., 2011, Taghizadeh-Sarabi et al., 2014, Stewart et al., 2014, Andersen et al., 2017], see [Lotte et al., 2007] for review.

We adopt a novel methodological approach for computing and evaluating SVM classifiers based on two approaches: 1) single-trial training and single-trial test classification, and 2) training on an averaged response of each of the 23 image categories for each subject (corresponding to 23 pseudo-trials per subject) and single-trial test classification. Furthermore, we open the black box and visualize which parts of the EEG signature are exploited by the SVM classifiers. In particular, we propose a method for computing sensitivity maps of EEG-based SVM classifiers based on a methodology originally proposed for fMRI [Rasmussen et al., 2011]. To evaluate effect sizes of sensitivity maps and event related potential (ERP) difference maps, we use a modified version of an NPAIRS resampling scheme [Strother et al., 2002]. Lastly, we investigate how the pseudo-trial classifier based on averaged category responses compares to the single-trial classifier in terms of prediction accuracy of novel subjects.

## 2. MATERIALS AND METHODS

### 2.1. Participants

A total of 15 healthy subjects with normal or corrected-to-normal vision (10 male, 5 female, mean age: 25, age range: 21-30), who gave written informed consent prior to the experiment, were recruited for the study. Participants reported no neurological or mental disorders. Non-invasive experiments on healthy subjects are exempt from ethical committee processing by Danish law [Den Nationale Videnskabsetiske Komité, 2014].

### 2.2. Stimuli

Stimuli consisted of 690 images from the Microsoft Common Objects in Context (MS COCO) dataset [Lin et al., 2014]. Images were selected from 23 semantic categories, with each category containing 30 images. Of the 23 categories, 10 categories contained animals and the remaining 13 categories contained inanimate items, such as food or man-made objects. Thus, each participant was exposed to 300 animate trials and 390 inanimate trials, resulting in a chance level of 56.5% for prediction of the larger, inanimate class. For categories and image labels used in the experiment, see Supplementary File 1. All images presented were unique and not repeated for the same subject throughout the experiment. The initial selection criteria were 1) image aspect ratio of 4:3, 2) only a single super- and subcategory per image, and 3) minimum 30 images within the category. Furthermore, we ensured that all 690 images had a relatively similar luminance and contrast to avoid the influence of low-level image features in the EEG signals. Thus, images within 77% of the brightness distribution and 87% of the contrast distribution were selected. Images that were highly distinct from standard MS COCO images were manually excluded (see Appendix A for exclusion criteria). Stimuli were presented using custom Python scripts built on PsychoPy2 software [Peirce, 2009].

### 2.3. Experimental Design

Participants were shown 23 blocks of trials composed of 30 images each. The order of categories and images within the categories was random for each subject. At the beginning of each category, a probe word denoting the category name was displayed for 5 s followed by the 30 images from the corresponding category. Each image was displayed for 1 s, set against a mid-grey background. Inter-stimuli intervals (ISI) of variable length were displayed between each image. The ISI length was randomly sampled according to a uniform distribution from a fixed list of ISI values between 1.85 s and 2.15 s in 50 ms intervals, ensuring an average ISI duration of 2 s. To minimize eye movements between trials, the ISI consisted of a white fixation cross superimposed on a mid-grey background in the center of the screen (Figure 1B).

Subjects viewed images on a computer monitor with a viewing distance of 57 cm. The size of stimuli was 4 × 3 degrees of visual angle. Duration of the experiment was 39.3 min, which included five 35 s breaks interspersed between the 23 blocks. Before the experimental start, participants underwent a familiarization phase with two blocks of reduced length (103 s).

### 2.4. EEG Data Collection

A user-friendly, portable EEG equipment, Enobio (Neuro-electrics) with 32 dry-electrode channels, was used for data acquisition. The EEG was electrically referenced using a CMS/DRL ear clip. The system recorded 24-bit EEG data with a sampling rate of 500 Hz, which was transmitted wirelessly using Wifi. LabRecorder was used for recording EEG signals. Lab Streaming Layer (LSL) was used to connect PsychoPy2 and LabRecorder for unified measurement of time series. The system was implemented on a Lenovo Legion Y520, and all recordings were performed in a normal office setting.

### 2.5. EEG Preprocessing

Among the 15 recordings, no participants were excluded during data preprocessing, as we would like to generalize our results to a broad range of experimental recordings. Preprocessing of the EEG was done using EEGLAB (sccn. ucsd.edu/eeglab). The EEG signal was bandpass filtered to 1-25 Hz using Finite Impulse Response filters, and down-sampled to 100 Hz. Artifact Subspace Reconstruction (ASR) [Mullen et al., 2015] was applied to reduce non-stationary high variance noise signals. Temporal trends in the EEG signals were investigated before and after ASR for each subject (Figures S2-S3). Generally, the time dependencies of the EEG signal were reduced by ASR. Channels which were removed by artifact rejection were interpolated from the remaining channels, and the data were subsequently re-referenced to an average reference. Epochs of 600 ms, 100 ms before and 500 ms after stimulus onset, similar to [Kaneshiro et al., 2015], were extracted for each trial.

A sampling drift of 100 ms throughout the entire experiment was observed for all subjects and was corrected for offline.

Since the signal-to-noise ratio varied across trials and participants, all signals were normalized to z-score values (i.e., each trial and averaged trials from each participant was transformed so that it had a mean value of 0 and a standard deviation of 1 across time samples and channels).

### 2.6. Support Vector Machines

Support vector machines (SVM) were implemented to classify the EEG data into two classes according to animate and inanimate trials. *y*_*i*_ ∈ {−1, 1} is the identifier of the category, and an observation is defined to be the EEG response in one epoch ([−100, 500] ms w.r.t. stimulus onset). SVMs allow adoption of a non-linear kernel function to transform input data into a high dimensional feature space, where it is possible to linearly separate data. The iterative learning process of the SVM will devise an optimal hyperplane with the maximal margin between each class in the high dimensional feature space. Thus, the maximum-margin hyperplane will form the decision boundary for distinguishing the brain response associated with animate and inanimate data [Saitta, 1995].

The SVM classifier is implemented by a non-linear projection of the observations **x**_**n**_ into a high-dimensional feature space 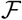.

Let 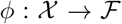 be a mapping from the input space 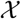 to 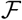. The weight vector **w** can be expressed as a linear combination of the training points 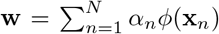 and the kernel trick is used to express the discriminant function as:

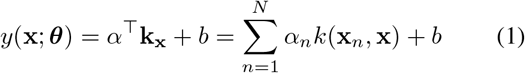

with the model now parametrized by the smaller set of parameters ***θ*** = {*α, b*} [Lautrup et al., 1994]. The Radial Basis Function (RBF) kernel allows for implementation of a non-linear decision boundary in the input space. The RBF kernel **k**_x_ holds the elements:

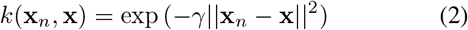

where *γ* is a tunable parameter.

Often it is desirable to allow a few misclassifications in the decision boundary in order to obtain a better generalization error. This trade-off is controlled by a tunable regularization parameter *c*.

Two overall types of SVM classifiers were implemented: 1) single-trial classifier, and 2) average category level classifier, denoted as pseudo-trial classifier based on terminology used in e.g. [Guggenmos et al., 2018]. Both classifiers decode supercategories, animate versus inanimate, and both classify between subjects. The single-trial classifier is trained on 690 trials for each subject included in the training set. The pseudotrial classifier averages the 30 trials within each of the 23 categories for each subject, such that the classifier is trained on 23 averaged, pseudo-trials for each subject included in the training set, instead of 690 trials.

The performance of the single-trial classifier was estimated using 14 participants as the training set, and the remaining participant was used as the test set (SVM parameters visualized in Figure S8). Cross-validation was performed on 10 parameter values in ranges *c* = [0.05; 10] and *γ* = [2.5 × 10^−7^; 5 × 10^−3^], thus cross-validating across 100 parameter combinations for each held out subject.

For a debiased estimate of the test accuracy, the single-trial classifier was trained on 13 subjects, with one participant held out for validation and another participant held out for testing, thus leaving out 2 subjects in each iteration. Fifteen classifiers were trained with different subjects held out in each iteration. An optimal parameter set of *c* and *γ* was estimated using participants 1-7 as validation subjects (mean parameter value), which was used to estimate the test accuracy for subjects 8-15 and vice versa. Thus, two sets of optimal parameters were found (Figure S10). Cross-validation was performed on 10 parameter values in ranges *c* = [0.25; 15] and *γ* = [5 × 10^−7^; 2.5 × 10^−2^], i.e. 100 combinations.

The pseudo-trial classifier is much faster to train and was built using a basic nested leave-one-subject-out cross-validation loop. In the outer loop, one subject was held out for testing while the remaining 14 subjects entered the inner loop. The inner loop was used to estimate the optimum *c* and *γ* parameters for the SVM classifier. The performance of the model was calculated based on the test set. Each subject served as test set once. A permutation test was performed to check for significance. For each left out test subject, the animacy labels were permuted and compared to the predicted labels. This was repeated 1000 times, and the accuracy scores of the permuted sets were compared against the accuracy score of the non-permuted set. The upper level of performance was estimated by choosing the parameters based on the test set. Cross-validation was performed on 10 parameter values in ranges *c* = [0.25; 15] and *γ* = [5 × 10^−7^; 2.5 × 10^−2^], i.e 100 combinations.

### 2.7. Sensitivity Map

To visualize the SVM RBF kernel, anapproach proposed by [Rasmussen et al., 2011] was adapted. The sensitivity map is computed as the derivative of the RBF kernel, c.f. Eq. (2)

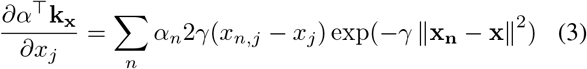

Pseudo-code for computing the sensitivity map across time samples and trials is found in Appendix B. A GitHub toolbox with Python implementation of sensitivity mapping is available: https://github.com/gretatuckute/DecodingSensitivityMapping.

### 2.8. Effect Size Evaluation

The NPAIRS (nonparametric prediction, activation, influence, and reproducibility resampling) framework [Strother et al., 2002] was implemented to evaluate effect sizes of the SVM sensitivity map and animate/inanimate ERP differences. The sensitivity map and the ERP difference map based on all subjects were thus scaled by the average difference of sub-sampled partitions.

The scaling was calculated based on *S* = 100 splits. In each split, two partitions of the data set were randomly selected without replacement. A partition consisted of 7 subjects, thus achieving two partitions of 7 subjects each (leaving a single, random subject out in each iteration).

For evaluation of the ERP difference map, a difference map was calculated for each partition (**M**_1_ and **M**_2_). Similarly, for evaluation of the sensitivity map, an SVM classifier was trained on each partition, and sensitivity maps were computed for both SVM classifiers (corresponding to **M**_1_ and **M**_2_ for the ERP difference map evaluation). The sensitivity map for the single-trial SVM classifier was computed based on optimal model parameters, while the sensitivity map of the pseudo-trial classifier was based on the mean parameters based on validation sets. The maps from the two partitions were contrasted and squared.

Across time samples (*t* = 1, …, *T*) and trials (*n* = 1, …, *N*) an average standard deviation of the average difference between partitions was calculated:

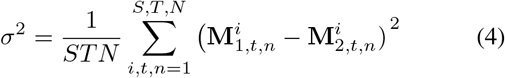

The full map, **M**_full_ (based on 15 subjects) was then divided by the standard deviation to produce the effect size

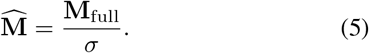

## 3. RESULTS

We classify the recorded EEG using SVM RBF models such that trials are labeled with the high-level category of their presented stimuli, i.e., either animate or inanimate. We first report results using a single-trial classifier followed by a pseudo-trial classifier using averaged category responses, and then apply the pseudo-trial classifier for prediction of single-trial EEG responses. Also, we report effect sizes of ERP difference maps and sensitivity maps for evaluation of both SVM classifiers.

### 3.1. Event Related Potential Analysis

After EEG data preprocessing and Artifact Subspace Reconstruction (ASR) (Section 2.5), we confirmed that our visual stimuli presentation elicited a visual evoked response. The ERPs for the trials of animate content and the trials of inanimate content are compared in Figure 2. The grand average ERPs across subjects (thick lines) are shown along with the average animate and inanimate ERPs of each subject.

It is indicated in Figure 2 which time samples were significant for the averaged selection of channels. A full map of significant time samples and channels can be seen in Figure S4. The significance level was controlled for multiple comparisons using the conservative Bonferroni correction.

The animate and inanimate ERPs were most different 310 ms after stimuli onset. This applied both for the selected channels in Figure 2 and in general, including frontal channels (Figure S4). The average scalp map for the two supercategories as well as the difference between them at 310 ms can be seen in Figure 2 in z-scored units.

Inspection of Figure 2 shows that visual stimuli presentation elicited a negative ERP component at 80-100 ms post-stimulus onset followed by a positive deflection at around 140 ms post-stimulus onset. A P300 subcomponent, P3a, was evident around 250 ms and a P3b component around 300 ms [Polich, 2007]. It is evident that the P3b component is more prominent for the animate category. The observed temporal ERP dynamics was comparable to prior ERP studies of the temporal dynamics of visual object processing [Cichy et al., 2014].

**Fig. 1.**
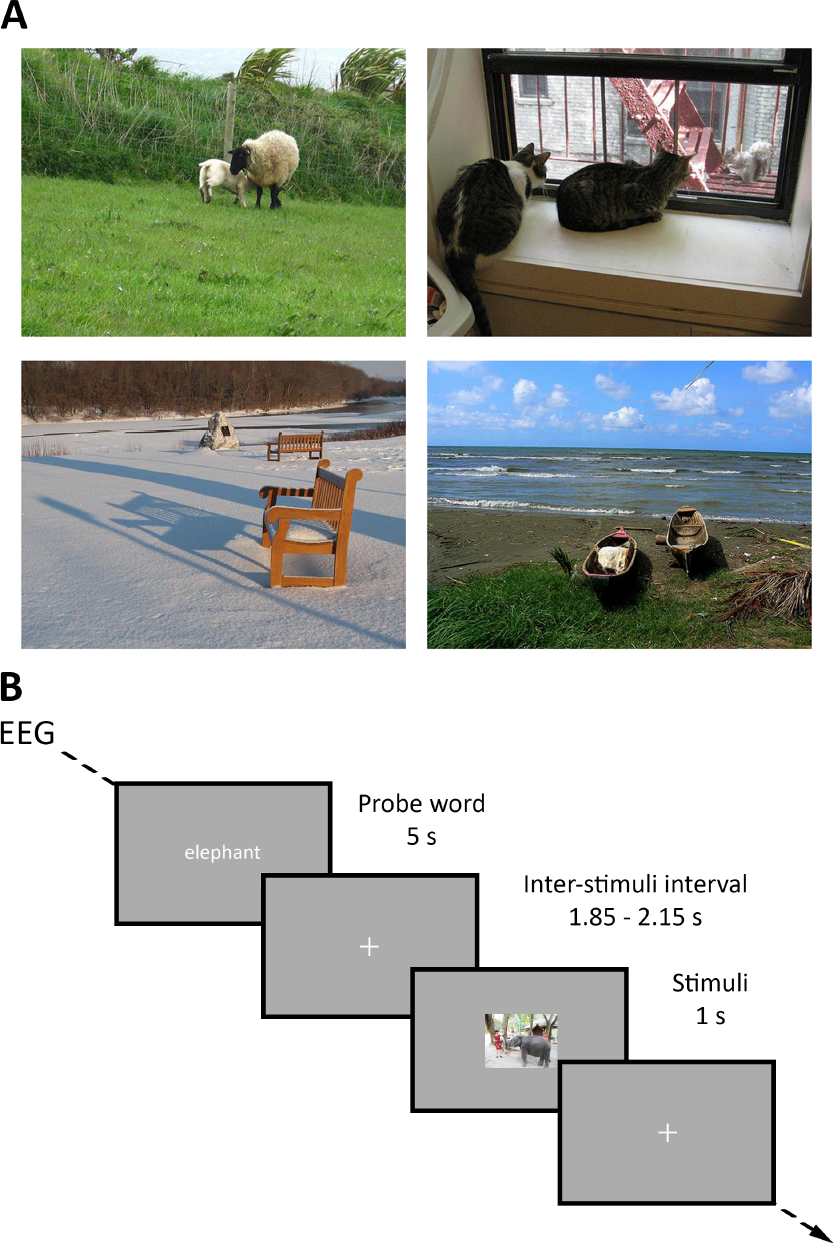
**A)** Example of the experimental visual stimuli. First row contains animate trials from “sheep” and “cat” categories, and second row contains inanimate trials from “bench” and “boat” categories. **B)** Experimental design of the visual stimuli presentation paradigm. The time-course of the events is shown. Participants were shown a probe word before each category, and jittered inter-stimuli intervals consisting of a fixation cross was added between stimuli presentation. The experiment consisted of 690 unique trials in total, 23 categories of 30 trials, ordered randomly (both category- and image-wise) for each subject.

Mean animate/inanimate ERP responses for each subject separately can be found in Figure S1.

### 3.2. Support Vector Machines

We sought to determine whether EEG data in our experiment can be automatically classified using SVM models. The Python toolbox Scikit-learn [Pedregosa et al., 2011] was used to implement RBF SVM models.

**Fig. 2.**
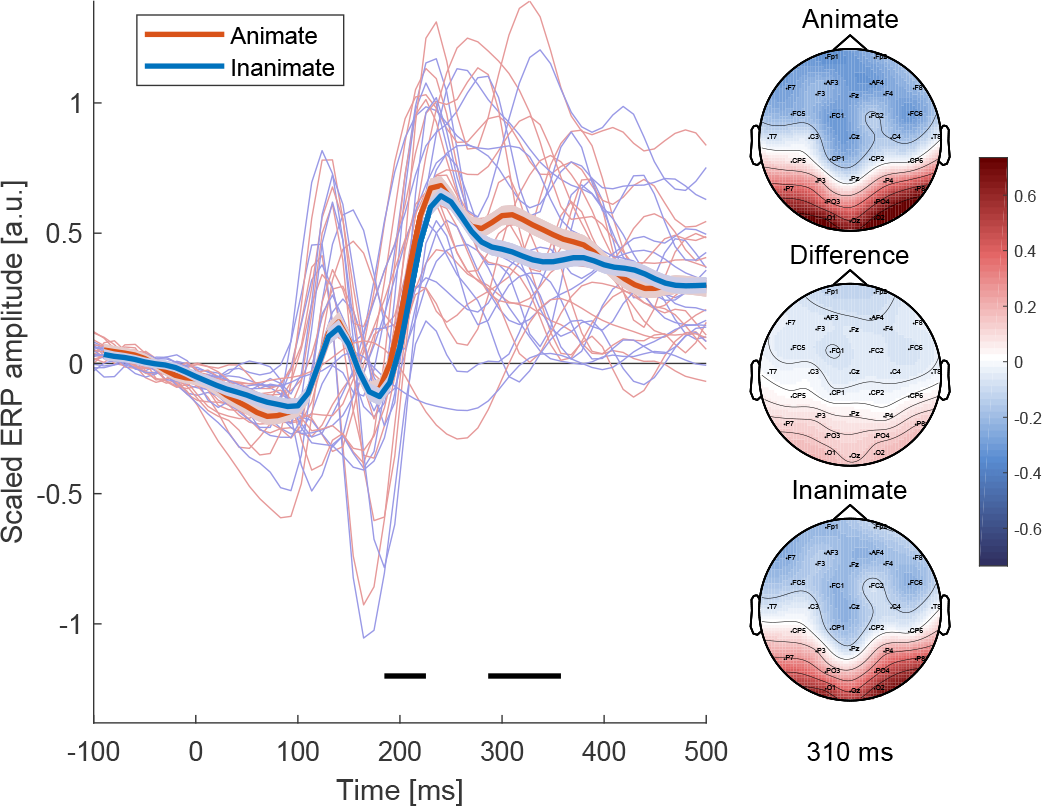
Average animate and inanimate ERPs across subjects (thick lines, and with standard errors around the mean) and for each subject (thin lines). ERP analysis was performed on the occipital/-parietal channels O1, O2, Oz, PO3, and PO4. The horizontal black lines indicate where the time samples are significant (paired t-test corrected for multiple comparisons, *α* = 0.05/60 time samples). Scalp maps are displayed for the animate/inanimate ERPs and difference thereof at 310 ms.

We specifically trained two different types of SVM classifiers, a single-trial, and a pseudo-trial classifier (averaged category responses), and assessed the classifiers’ accuracy on labeling EEG data in a leave-one-subject-out approach.

SVMs are regarded efficient tools for high-dimensional binary as well as non-linear classification tasks, but their ultimate classification performance depends heavily upon the selection of appropriate parameters of *c* and *γ* [Bishop, 2006]. Parameters for the upper level of performance for the single-trial classifier were found using cross-validation in a leave-one-subject-out approach, resulting in a penalty parameter *c* = 1.5 and *γ* = 5 × 10^−5^ based on the optimum mean parameters across test subjects (Figure S9). From Figure S8 it is evident that the optimum parameters were different for each subject, underlining inter-subject variability in the EEG responses.

To reduce bias of the performance estimate of the single-trial classifier, parameters were selected based on two validation partitions, resulting in *c* = 0.5 and *γ* = 5 × 10^−4^ for the first validation set, and *c* = 1.5 and *γ* = 5 × 10^−5^ for the second validation set (Figure S10).

The pseudo-trial classifier also showed inter-subject variability with respect to the model parameters, see Figures S5-S7. The classifier had an average penalty parameter of *c* = 7.2, and an average *γ* = 3.7 × 10^−4^ when based on the validation sets. The average optimum parameters when based on test sets with averaged categories and single-trials were in the same range, with *c* = 4.4 and *γ* = 3.0 × 10^−4^ and *c* = 6.7 and *γ* = 2.2 × 10^−4^, respectively.

**Fig. 3.**
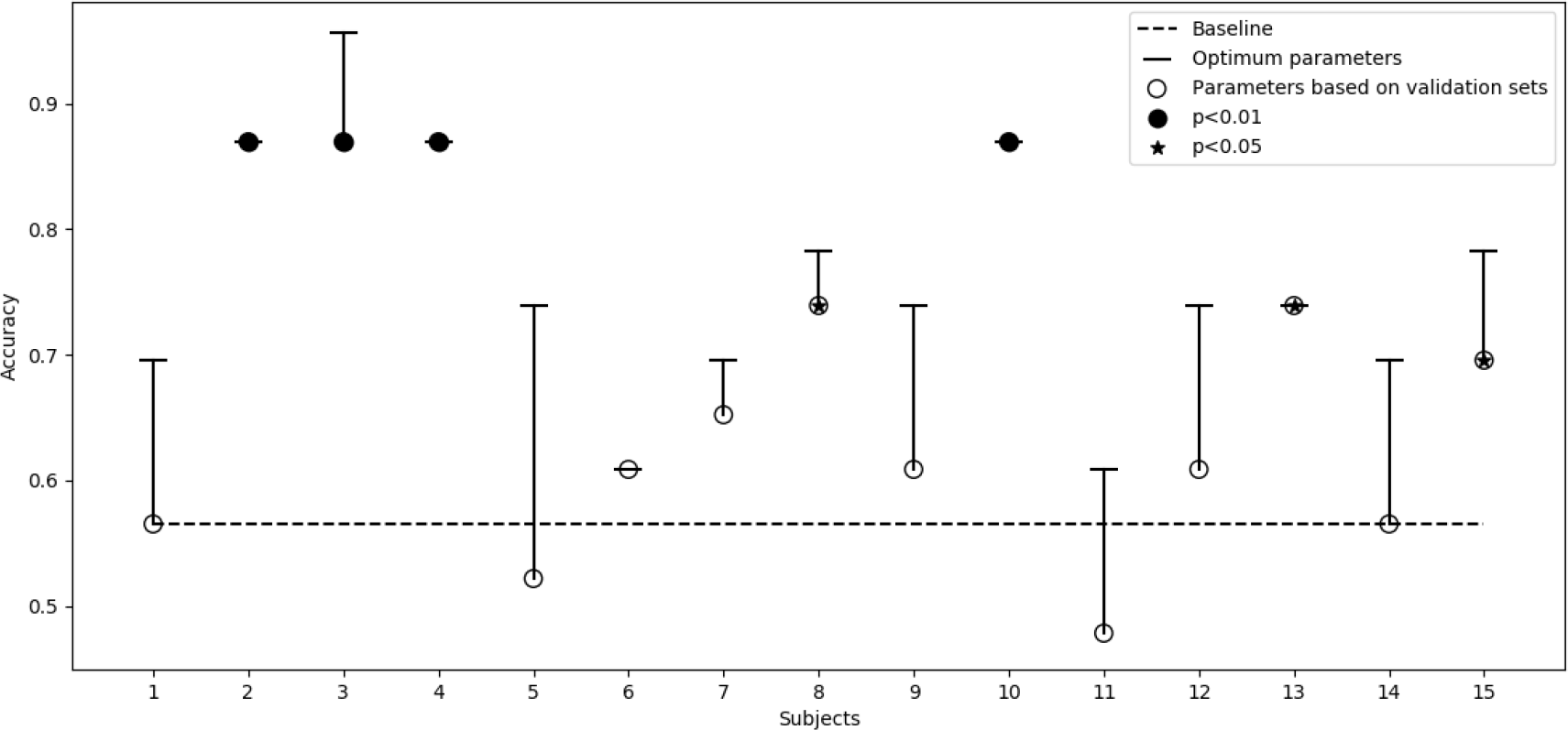
Test accuracies for the SVM pseudo-trial classifier trained on average categories and tested on average categories. The x-axis refers to the hold-out test subject. Chance level prediction accuracy is 0.565 (dashed line). Significance estimated by permutation testing (1000 iterations).

**Fig. 4.**
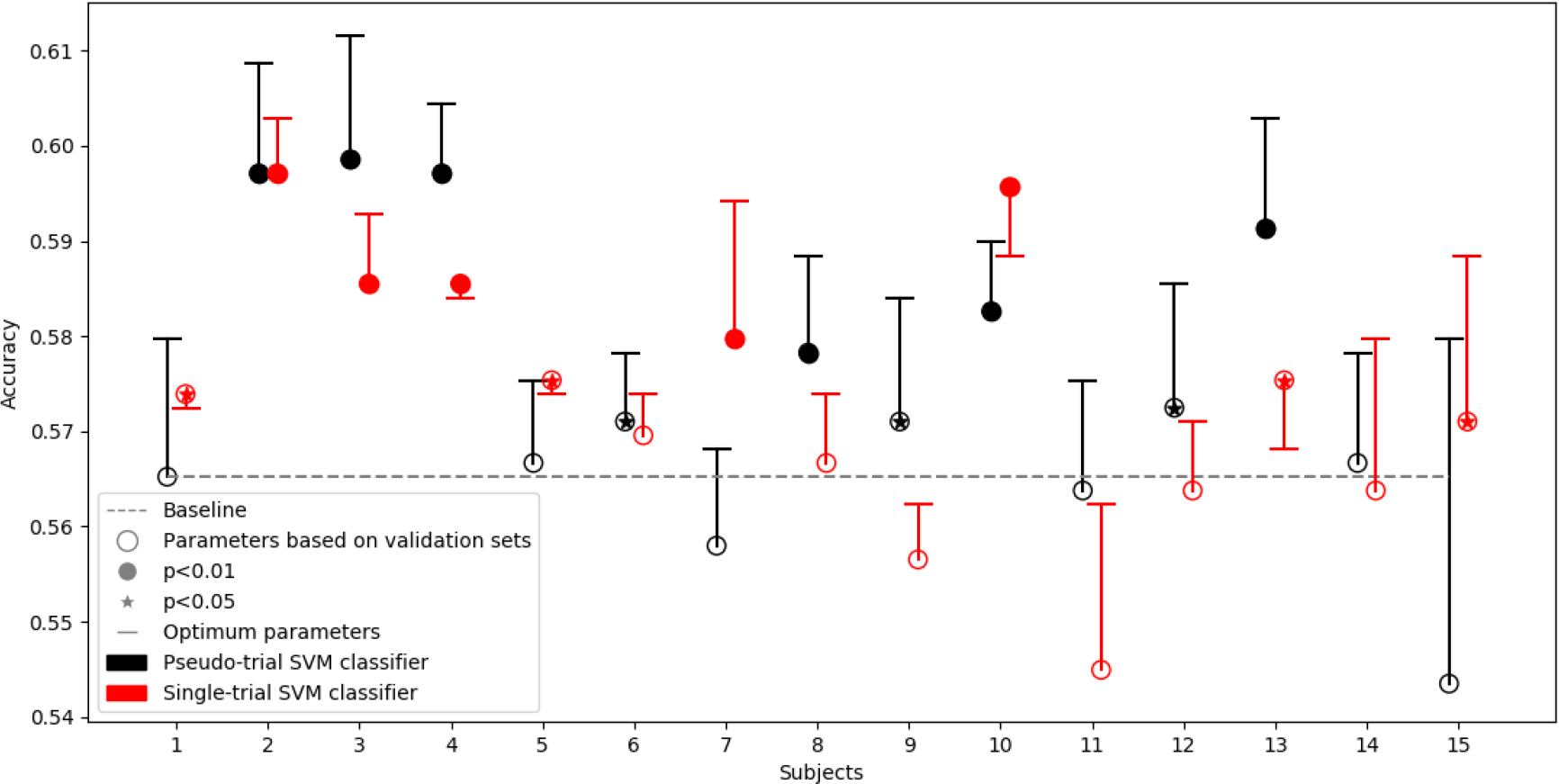
Test accuracies for a classifier trained on pseudo-trials (averaged categories) (black) or trained on single-trials (red) and tested on single-trials. The x-axis refers to the hold-out test subject. Chance level prediction accuracy is 0.565 (dashed line). Note: In some cases the “optimum parameters” are not found to be optimum, which can be explained by different training phases of the two single-trial classifiers. The classifier based on validation sets was trained on 13 subjects while the classifier with parameters based on the test set was trained on 14 subjects. For 5 out of 15 subjects the classifier based on 13 subjects was able to obtain higher accuracies.

**Fig. 5.**
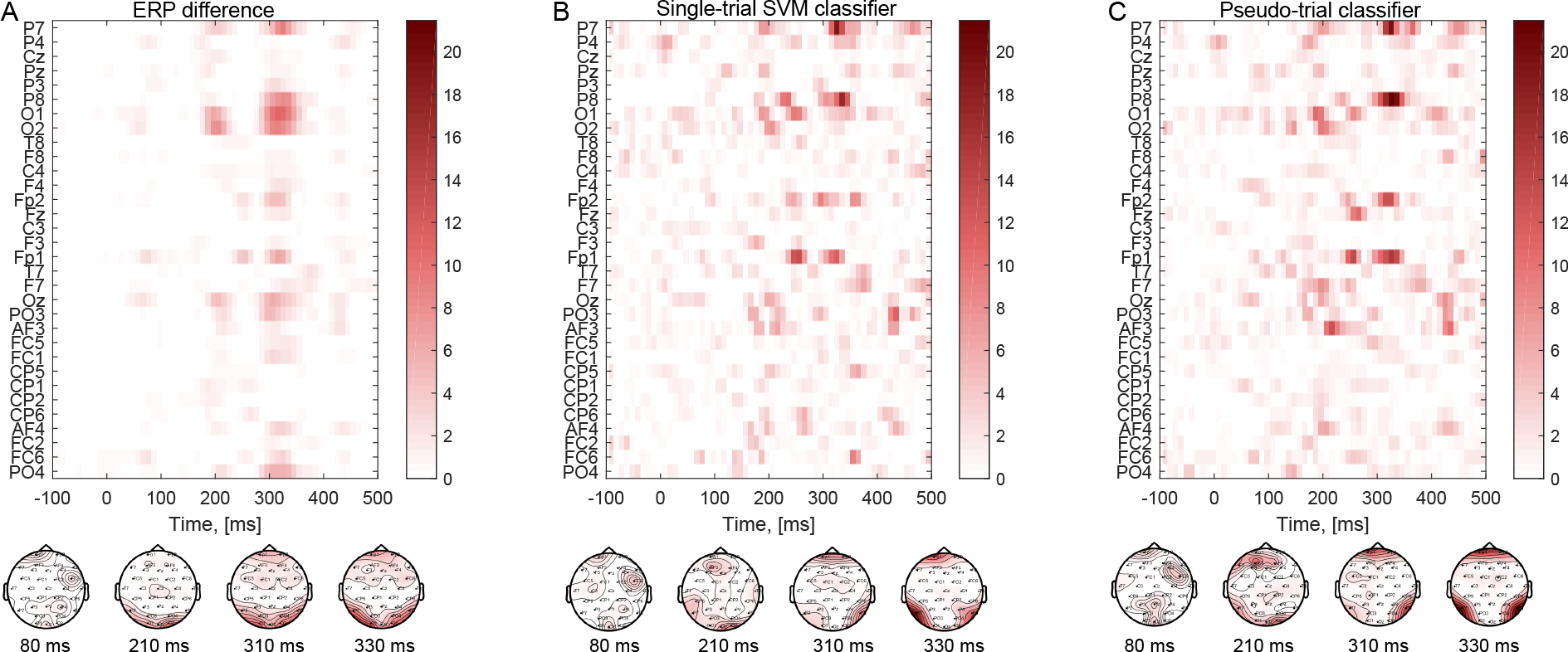
**A)** Effect sizes for animate/inanimate ERP difference map. **B)** and **C)** Effect sizes for the sensitivity maps of the single-trial and pseudo-trial SVM classifiers. Effect sizes were computed based on 100 NPAIRS resampling splits.

Figures 3-4 show the SVM classification performances using the two types of classifiers. Based on the leave-one-subject-out classification, we note the large variability of single subject performance. While different performances are obtained using the single-trial and pseudo-trial classifiers on single-trial test sets, the overall accuracies are similar (p=0.82, paired t-test), with an average of 0.574 and 0.575, respectively (Figure 4). Thus, the pseudo-trial classifier performs on par with the single-trial classifier in the prediction of single-trial test subjects.

A standard error of the mean of 0.01 was found for both the debiased performance measure of the single-trial classifier and for the unbiased single-trial classifier (corrected for the leave-one-subject-out approach [Efron and Tibshirani, 1994])

### 3.3. Event Related Potential Difference Map and Sensitivity Map

We investigated the raw ERP difference map between animate and inanimate categories, as well as the sensitivity maps for the single-trial and pseudo-trial SVM classifiers. The sensitivity map reveals EEG time points and channels that are of relevance to the SVM decoding classifiers (Figure 5).

For map effect size evaluation we implement an NPAIRS resampling scheme [Strother et al., 2002]. In this cross-validation framework, the data were split into two partitions of equal size (7 subjects in each partition randomly selected without replacement). This procedure was repeated 100 times to obtain standard errors of the maps for computing effect sizes (Section 2.8).

Figure 5A displays the effect sizes of the raw ERP difference map between the animate and inanimate categories, while Figure 5B and 5C displays effect sizes of sensitivity maps for the single-trial and pseudo-trial classifiers, respectively. Scalp maps show the spatial information exploited by the classifiers at different time points.

From inspection of Figure 5 it is evident that occipital and parietal channels (O1, O2, P7, P8) were relevant for SVM classification at time points comparable to the ERP difference map. Frontal channels (Fp1, Fp2) were exploited by both SVM classifiers, but to a larger extent by the pseudo-trial classifier (Figure 5C). Furthermore, the pseudo-trial classifier exploited a larger proportion of earlier time points compared to the single-trial classifier. The sensitivity maps for the singletrial and pseudo-trial classifiers suggest that despite the difference in number and type of trials, the classifiers are similar.

## 4. DISCUSSION

In the current work, we approach the challenges of EEG-based decoding: non-laboratory settings, user-friendly wireless EEG acquisition equipment with dry electrodes, natural stimuli, no repetition of experimental stimuli trials, and no exclusion of participants. Thus, our work is centered around 1) ecological validity and portability, and 2) generalizability. The potential benefits of mitigating these challenges is to study the brain dynamics in natural settings and for applications in real-life scenarios.

Our motivation for working with a portable, dry-electrode EEG system is to increase the EEG usability in terms of affordability, mobility, and ease of maintenance. These factors are crucial in applied contexts in everyday settings, such as the development of real-time EEG neurofeedback systems. It has recently been demonstrated that commercial-grade EEG equipment compares to high-grade equipment in laboratory settings in terms of neural reliability as quantified by inter-subject correlation [Poulsen et al., 2017]. Furthermore, a systematic comparison between a wireless dry EEG system and a conventional laboratory-based wet EEG system shows similar performance in terms of signal quality [Kam et al., 2019].

We aim to increase the generalization ability of our decoding models. To do so, we evaluate decoding ability in an inter-subject design, i.e., leave-one-subject-out approach [Kjems et al., 2002]. Prior studies in EEG-based decoding, in particular for BCIs, have focused on building classifiers to decode subject-specific brain patterns, see [Cecotti, 2011] for review. Inter-subject generalized BCI has the advantage of saving time in BCI sessions, and several research groups have made effort to develop inter-subject generalized BCI systems for decoding of motor imagery-related EEG [Ray et al., 2015, Halme and Parkkonen, 2018]. Successful inter-subject classification requires extraction of globally relevant signal features from each training subject [Kjems et al., 2002]. In the current work, we take a step towards increasing generalizability by building inter-subject EEG-based decoding models.

Our ultimate goal is to decode actual semantic differences between natural categories; thus we perform low-level visual feature standardization of experimental trials prior to the experiment, investigate time dependency of the EEG response throughout the experiment, and perform ASR to reduce this dependency (Section 2.5). Moreover, the stimuli in our experimental paradigm consisted of complex everyday scenes and non-iconic views of objects [Lin et al., 2014]. Animate and inanimate images were similar in composition, i.e., an object or animal in its natural surroundings (Figure 1A).

### 4.1. Data Preprocessing of Temporal Trends

There will naturally be continuous variations in EEG recordings over time. Since our experimental paradigm lasted approximately 40 minutes, we investigated temporal trends in the EEG data (Figures S2-S3) and perform artifact subspace reconstruction (ASR) [Mullen et al., 2015] to reduce confounding temporal trends in further analyses. The unwanted non-stationarity of the EEG signal arises from electrodes gradually losing or gaining connection to the scalp, an increasing tension of facial muscles or other artifactual currents [Delorme et al., 2007, Rowan and Tolunsky, 2003]. If the data are epoched, the drift may misleadingly appear as a pattern reproducible over trials, a tendency that may be further reinforced by component analysis techniques that emphasize repeatable components [de Cheveigné and Arzounian, 2018]. Slow linear drifts can be removed by employing high-pass filters, however more complicated temporal effects are harder to remove. Furthermore, employing high-pass filters may risk introducing new artifacts. As an alternative recent studies suggest performing robust detrending, where the trend of each channel is determined and then regressed out [de Cheveigné and Arzounian, 2018, Driel et al., 2019]. We observe that by employing ASR the time dependency was reduced for most subjects (Figures S2-S3). However, it would be interesting to investigate more complex detrending algorithms to also make sure that high-pass filtering is not impairing our results.

### 4.2. Event Related Potential Analysis

Previous work on visual stimuli decoding demonstrate semantic category specificity at both early (~ 150 ms) and late (~ 400 ms) intervals of the visually evoked potential [Rousselet et al., 2004, Rousselet et al., 2007]. ERP studies indicate that category-attribute interactions (natural/non-natural) emerge as early as 116 ms after stimulus onset over frontocentral scalp regions, and at 150 and 200 ms after stimulus onset over occipitoparietal scalp regions [Hoenig et al., 2008]. Kaneshiro et al., 2015, demonstrate that the first 500 ms of single-trial EEG responses contain information for successful category decoding between human faces and objects, and above chance object classification as early as 48-128 ms after stimulus onset [Kaneshiro et al., 2015]. For animate versus inanimate images, ERP differences have been demonstrated detectable within 150 ms of presentation [Thorpe et al., 1996]. However, there appears to be uncertainty whether these early ERP differences represent low-level visual stimuli or actual high-level differences. We observe the major difference between animate/inanimate ERPs around 210 ms and 320 ms (Figure 2 and S4). Akin to our results, Carlson et al., 2013 found that high-level categories (animacy) were maximally decodable around 240 ms from MEG recordings [Carlson et al., 2013]. Lastly, we observe that ERP signatures were highly variable among subjects (comparable to [Simanova et al., 2010]), which challenges the inter-subject model generalizability with our sample size of 15 subjects.

### 4.3. Support Vector Machine Classification

In this study, we adopted RBF kernel SVM classifiers to classify between animate/inanimate natural visual stimuli in a leave-one-subject-out approach. SVM classifiers have previously been implemented for EEG-based decoding. SVM in combination with independent component analysis data processing has been used to classify whether a visual object is present or absent from EEG [Stewart et al., 2014]. Zafar et al., 2017, propose a hybrid algorithm using convolutional neural networks for feature extraction and likelihood-ratio-based score fusion for prediction of brain activity from EEG [Zafar et al., 2017]. Taghizadeh-Sarabi et al., 2015, extract wavelet features from EEG, and selected features are classified using a “one-against-one” SVM multiclass classifier with optimum SVM parameters set separately for each subject [Taghizadeh-Sarabi et al., 2014].

We implemented single-trial and pseudo-trial (i.e. averaged categories) SVM classifiers, and found very similar performance of the single-trial and pseudo-trial classifiers for prediction of single-trial subjects (Figure 4). As the pseudotrial classifier is significantly faster to train, a full nested cross-validation scheme was feasible. The fact that the two classifiers have similar performance indicates that the reduced sample size in the pseudo-trial classifier is offset by the better signal-to-noise ratio of averaged trials. The fast training of the pseudo-trial classifier allows for parameter optimization and unbiased performance evaluation.

Based on the leave-one-subject-out classification performance (Figures 3-4), it is evident that there is a difference in how well the classifier generalizes across subjects, which partly is due to the diversity of ERP signatures across subjects (Figure S1). For some subjects, low accuracy is caused by a parameter mismatch between trials belonging to that subject and its validation sets. For other subjects, the SVM model is not capable of capturing their data even when parameters are based on that subject, due to poor signal-to-noise level. Furthermore, inter-subject generalizability in EEG is complicated by multiple factors: The signal-to-noise ratio at each electrode is affected by the contact to the scalp which is influenced by local differences in skin condition and hair, the spatial location of electrodes relative to underlying cortex will vary according to anatomical head differences, and there may be individual differences in functional localization across participants.

Both SVM classifiers utilized a relatively large number of support vectors. The single-trial SVM classifier used for computing the sensitivity map had model coefficients *α* = −1.5, …, 1.5, where 1204 *α* values out of 10350 were equal to 0 (9146 support vectors). The pseudo-trial classifier had model coefficients in the range *α* = −7.2, …, 7.2, and 46 out of 345 coefficients were zero (299 support vectors). The high number of obtained support vectors indicates a poor EEG signal-to-noise ratio and the complexity of the classification problem [Saitta, 1995].

### 4.4. Sensitivity Mapping

In the current work, we ask which parts of the EEG signatures are being exploited by the SVM decoding classifiers. We investigated the probabilistic sensitivity map for single-trial and pseudo-trial SVM classifiers based on a binary classification task. We identified spatial and temporal regions where discriminative information resides, and found these EEG features comparable to the difference map between raw ERP responses for animate and inanimate trials. We observe the most prominent difference in animate/inanimate ERPs around 210 ms and 320 ms (Figure 2 and S4), and these time points are also exploited by the SVM classifiers to a large extent (Figure 5).

The sensitivity maps for both SVM classifiers reveal that the occipital/-parietal channels where visual stimuli is known to be processed [Simanova et al., 2010, Kaneshiro et al., 2015] are major channels of interest in the classification task. Furthermore, we note that Fp1 and Fp2 channels are also important in the constructed classifiers (Figure 5). These two frontal channels also display significant differences in animate/inanimate ERPs across all subjects (Figure S4), which might be explained by a difference in eye movements depending on semantic category. Some studies report that frontal cortex activation is involved in distinguishing between visual stimuli [Wang et al., 2012], and it has been proposed that frontal activation during visual processing is a result of the attentional and anticipatory state of the subject [Foxe and Simpson, 2002]. However, it also possible that the frontal channels explain the noise in the informative channels [Blankertz et al., 2011].

Based on the similarity between the sensitivity maps for single-trial and pseudo-trial classifiers (Figure 5), we conclude that these classifiers exploit the same EEG features to a large extent. We therefore investigated whether the pseudotrial classifier is able to predict on single-trial test subjects. We demonstrate that classifiers trained on averaged pseudotrials perform on par with classifiers trained on a large set of noisy single-trial samples (Figure 4).

### 4.5. Conclusion

We investigate scalp EEG recorded with a portable 32 dry-electrode EEG equipment from healthy subjects under natural stimuli. We accomplish unbiased decoding of singletrial EEG using SVM models trained on de-noised (averaged) pseudo-trials, thus facilitating fast classifier training, parameter optimization and unbiased performance evaluation. The SVM classifiers were evaluated in a inter-subject approach, thus probing generalizability across participants. We propose a novel methodology for evaluating and computing sensitivity maps for EEG-based SVM classifiers, allowing for visualization of discriminative SVM classifier information. We implement an NPAIRS resampling scheme to compute sensitivity map effect sizes, and demonstrate high similarity between sensitivity map effect sizes of classifiers trained on small samples of de-noised, averaged data (pseudo-trial) and large samples of noisy data (single-trial). Finally, by linking temporal and spatial features of EEG to training of SVM classifiers, we take an essential step in understanding how machine learning techniques exploit neural signals.

## 5. OVERVIEW SUPPLEMENTARY MATERIAL

Appendix A: Manual exclusion criteria for image selection.

Appendix B: Sensitivity map Python pseudo-code.

Supplementary file 1: Image IDs, supercategories and categories for all images used in the experiment from Microsoft Common Objects in Context (MS COCO) image database.

Figures S1-S10 contain supplementary material, and are used for reference in the main manuscript.

## 6. DATA AVAILABILITY

Code available: https://github.com/gretatuckute/DecodingSensitivityMapping.

## 9. AUTHORS’ CONTRIBUTIONS

G.T, N.P and L.K.H. designed research; G.T and N.P acquired data; D.S, G.T and N.P performed initial data analyses; G.T., S.T.H and L.K.H performed research; G.T, S.T.H and L.K.H wrote the paper.

## 10. APPENDIX

## Appendix A

Manual exclusion criteria for MS COCO images [Lin et al., 2014] for the experimental paradigm:

- Object unidentifiable
- Object not correctly categorized
- Different object profoundly more in focus
- Color scale manipulation
- Frame or text overlay on image
- Distorted photograph angle
- Inappropriate image

## Appendix B

The following piece of pseudo-code shows how to compute the sensitivity map for an SVM classifier with an RBF kernel across all trials using Python and NumPy (np).

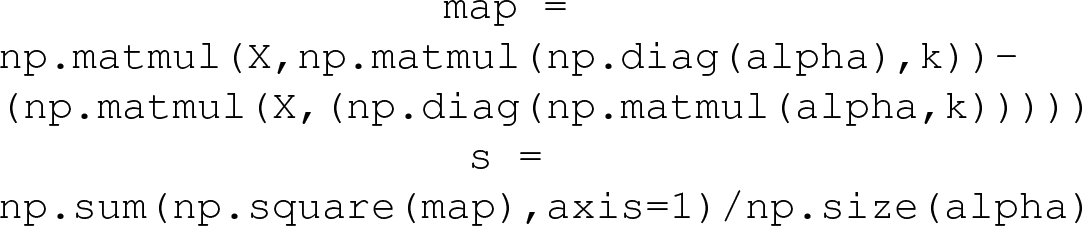

`k` denotes the (*N × N*) RBF training kernel matrix from equation 2, with *N* as the number of training examples. `alpha` denotes a (1 × *N*) vector with model coefficients. X denotes a (*P × N*) matrix with training examples in columns. s is a (*P* × 1) vector with estimates of channel sensitivities for each time point, which can be re-sized into a matrix of size [no. channels, no. time points] for EEG-based sensitivity map visualization.

## Supplementary materials

**Fig. S1.**
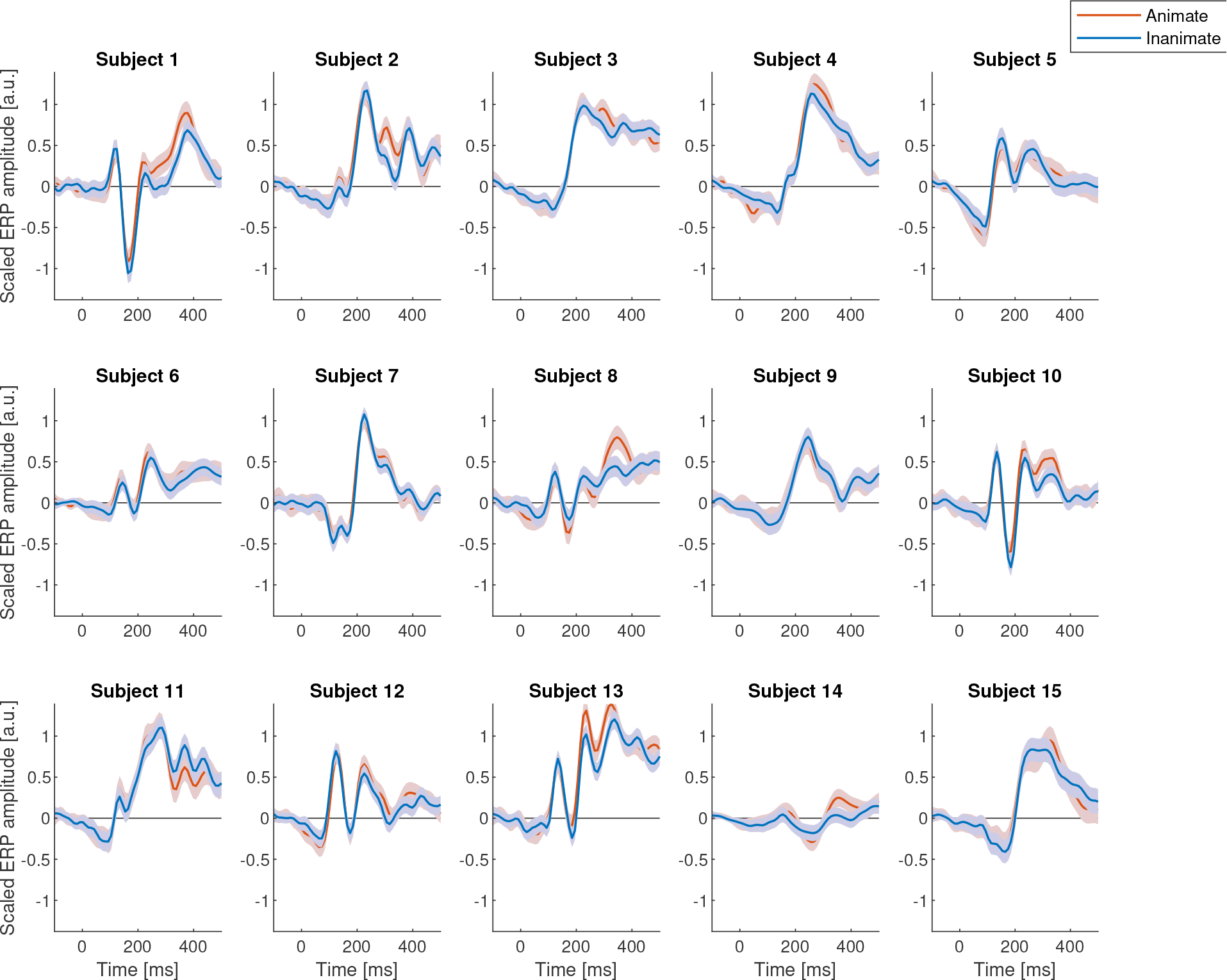
Animate and inanimate ERPs for each subject separately with two standard errors around the mean.

**Fig. S2.**
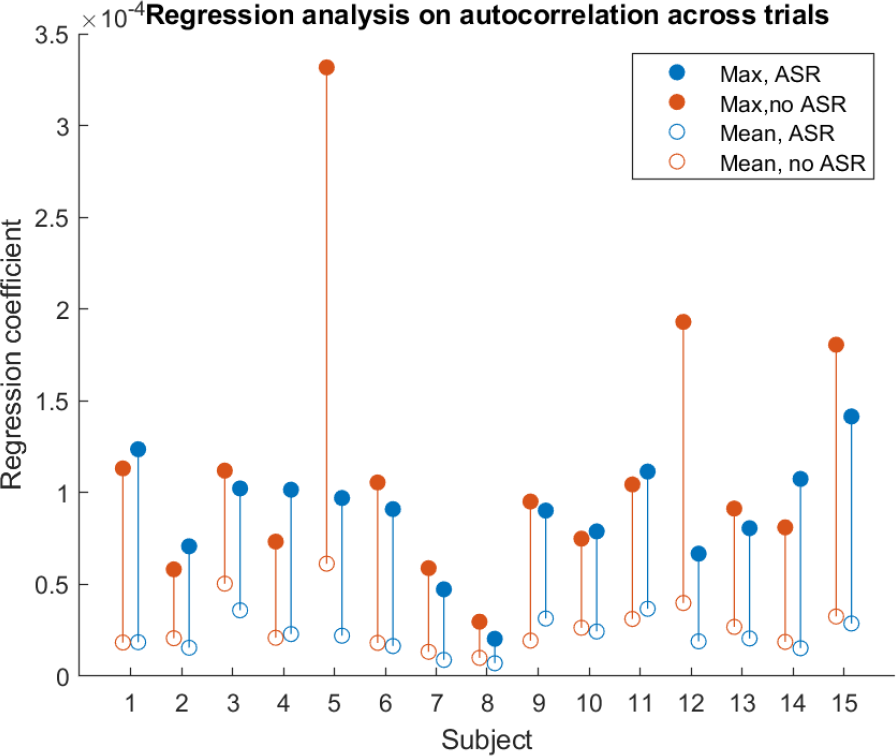
Time dependency as quantified by autocorrelation. A high regression coefficient means that a channel (or an average of all channels) had an autocorrelation which increased or decreased linearly with time lags, and is thus indicative of high time dependency.

**Fig. S3.**
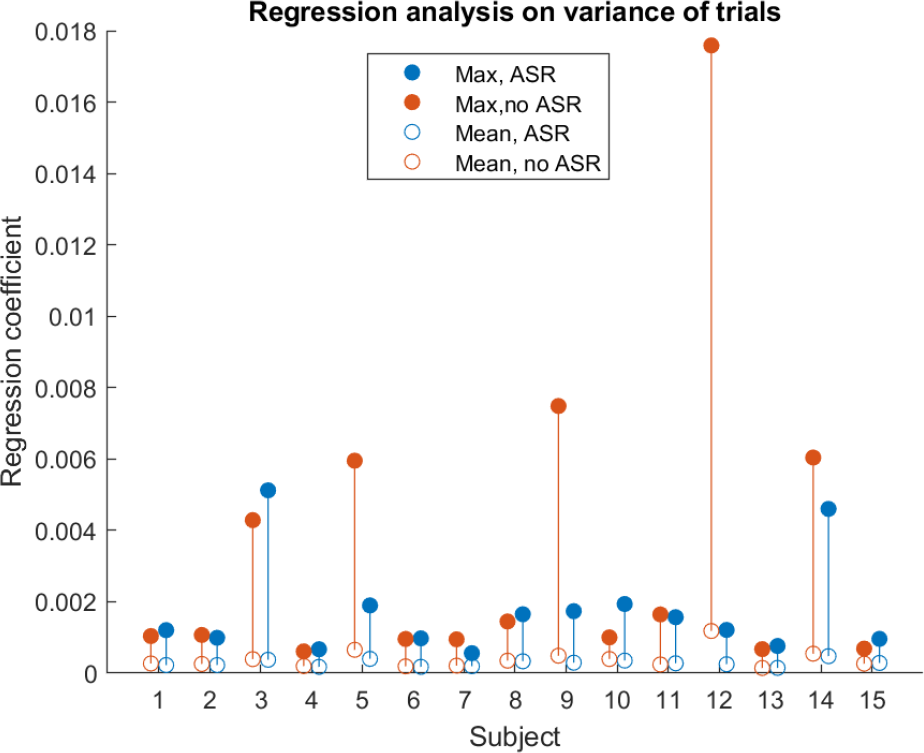
Time dependency as quantified by variance of trials. A high regression coefficient means that a channel (or an average of all channels) had a trial variance which increased or decreased linearly with trial number, and is thus indicative of high time dependency.

**Fig. S4.**
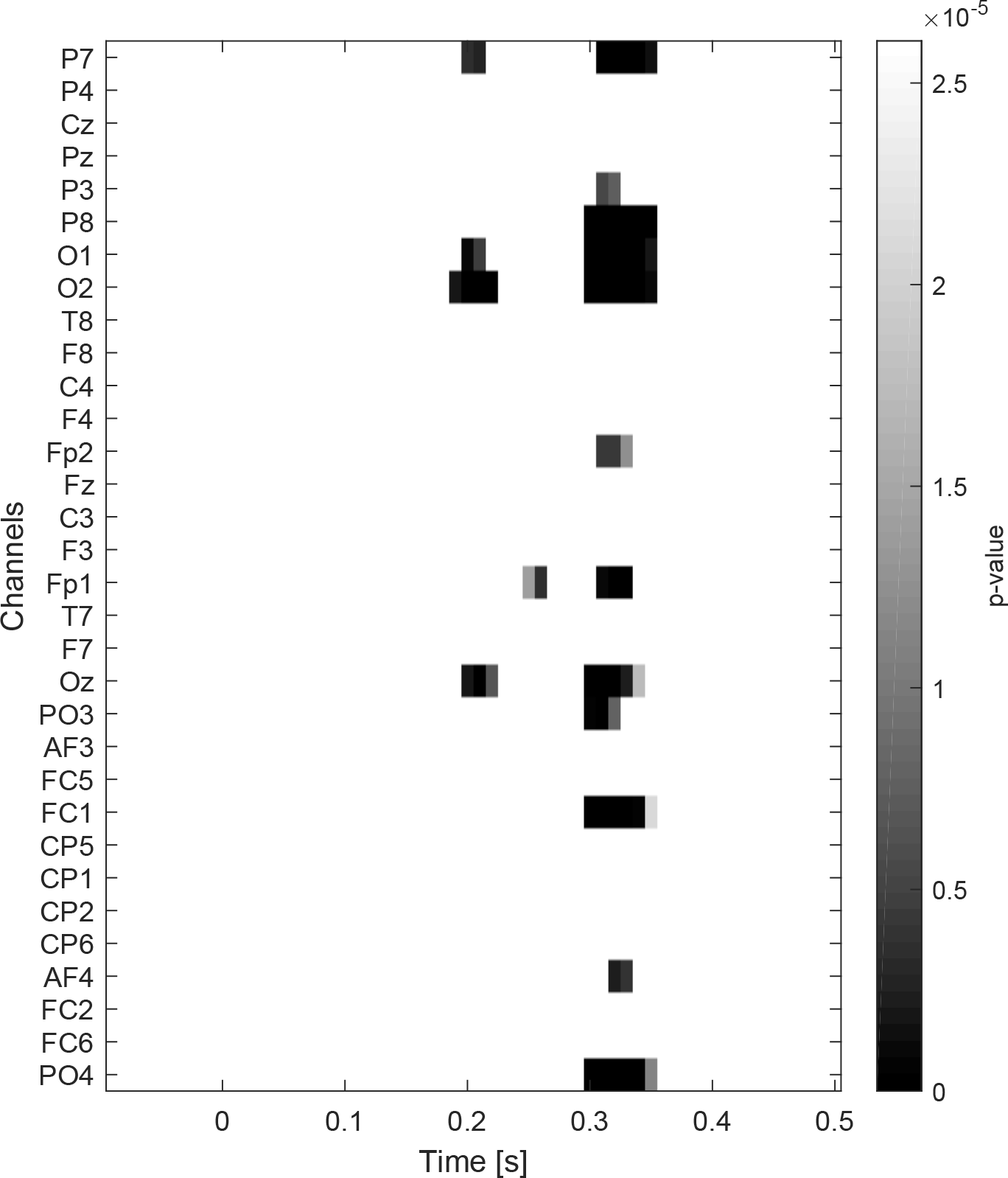
Significant ERP differences between animate and inanimate trials (all subjects). Significance tested using a paired t-test. Figure thresholded at *α* = 0.05/(60 · 32), i.e. Bonferroni corrected for multiple comparisons.

**Fig. S5.**
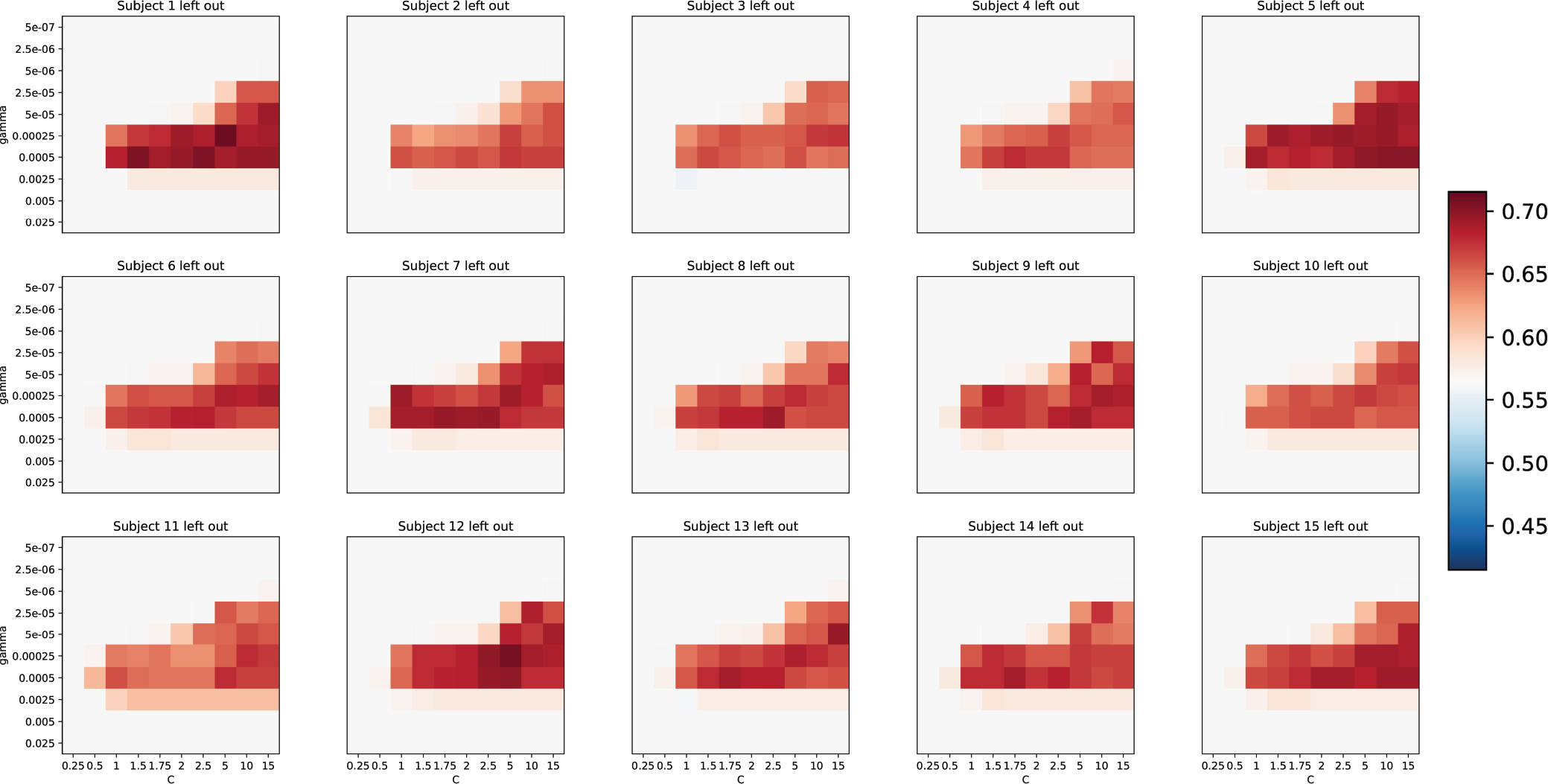
Validation accuracies (mean over validation sets) for the pseudo-trial classifier. *c* values are displayed on the x-axis, and consisted of values: [0.25, 0.5, 1, 1.5, 1.75, 2, 2.5, 5, 10, 15]. *γ* values are displayed on the y-axis, and consisted of values: [5 × 10^−7^, 2.5 × 10^−6^, 5 × 10^−6^, 2.5 × 10^−5^, 5 × 10^−5^, 2.5 × 10^−4^, 5 × 10^−4^, 2.5 × 10^−3^, 5 × 10^−3^, 2.5 × 10^−2^]. Same scaling for all subjects.

**Fig. S6.**
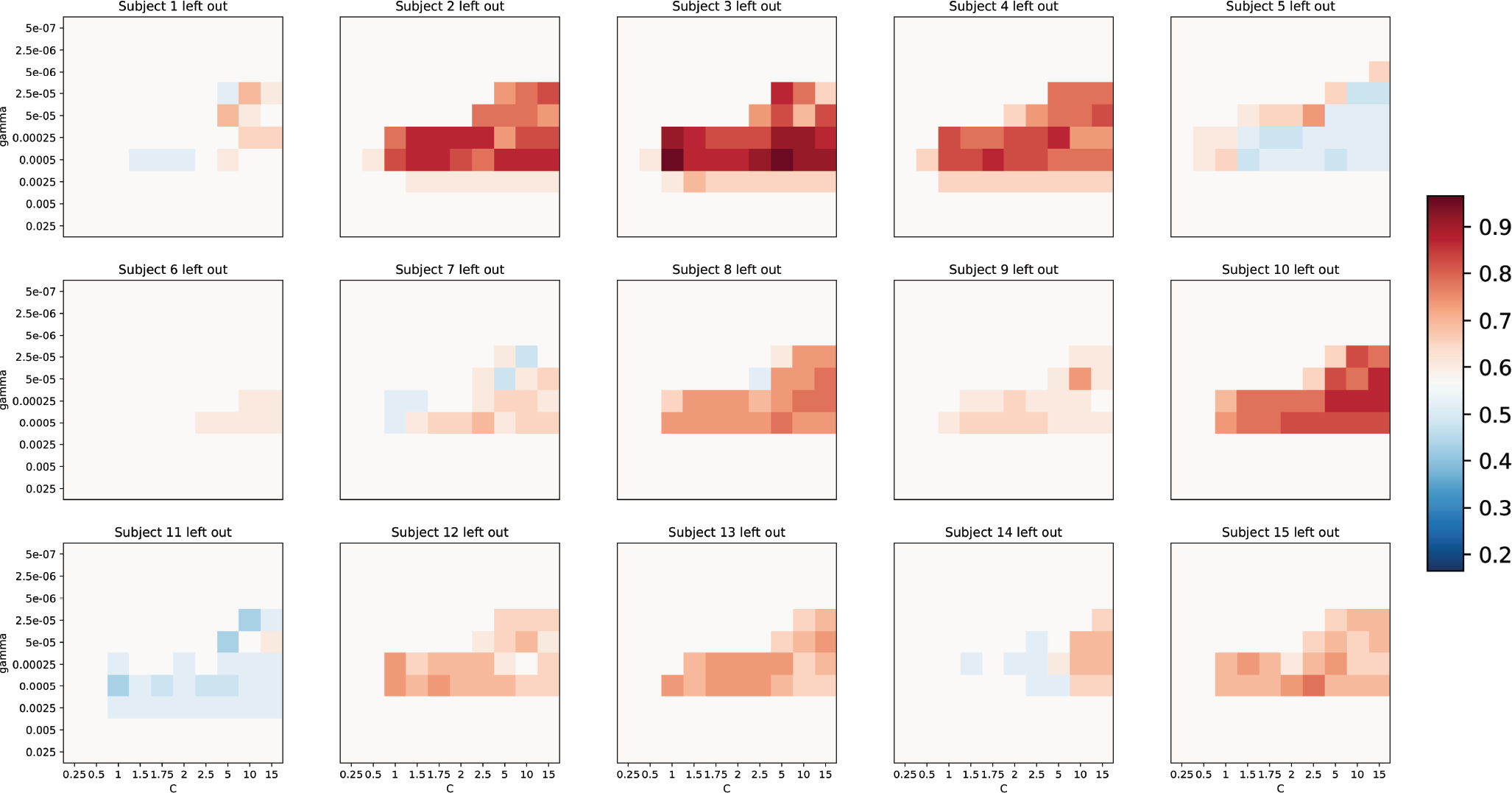
Test accuracies for pseudo-trial classifier. Tested on the pseudo-trials (averaged categories) of the withheld subject. Same cross-validation parameter values as in Figure S5.

**Fig. S7.**
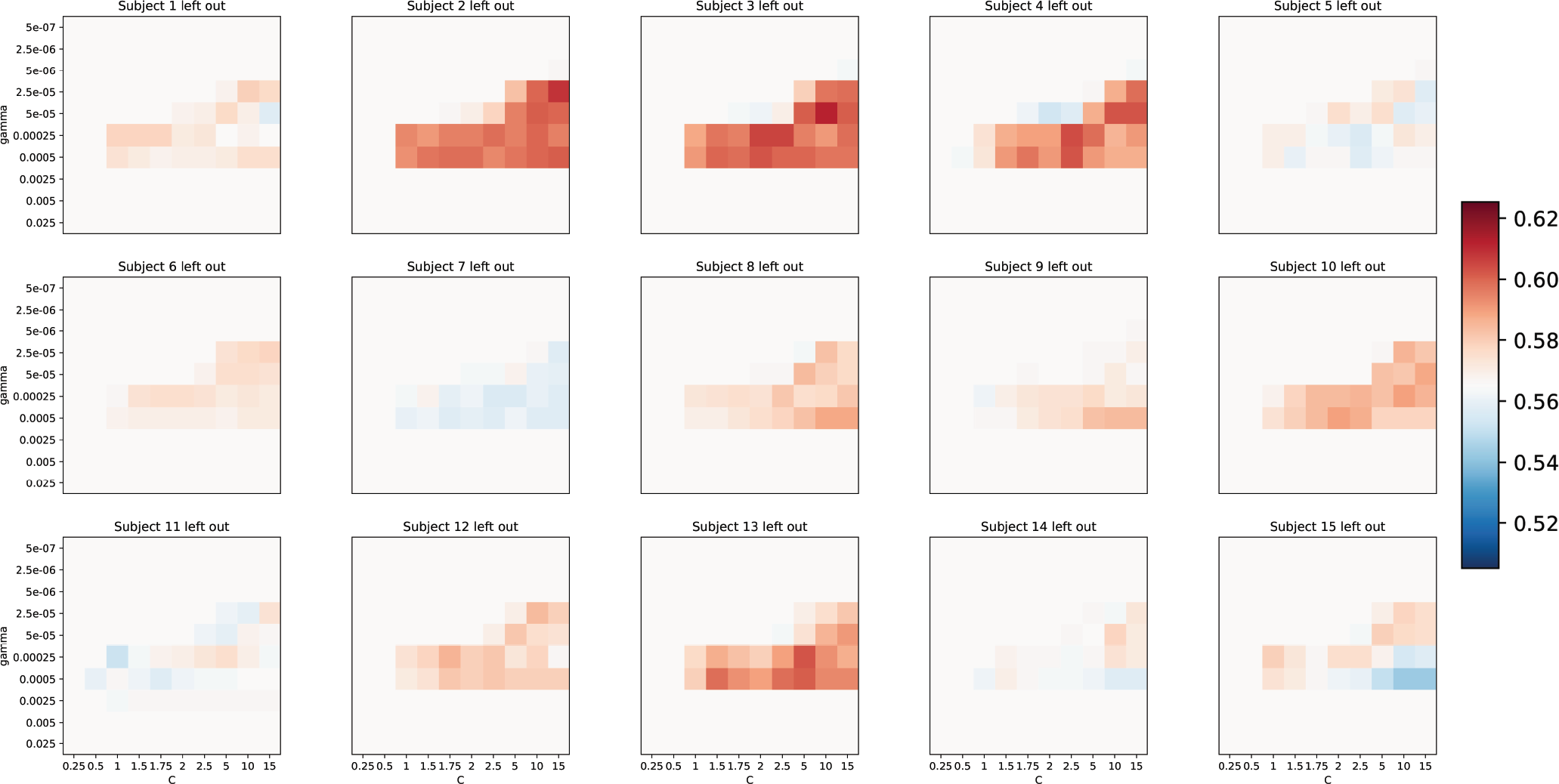
Test accuracies for pseudo-trial classifier. Tested on the pseudo-trials (averaged categories) of the withheld subject. Same cross-validation parameter values as in Figure S5.

**Fig. S8.**
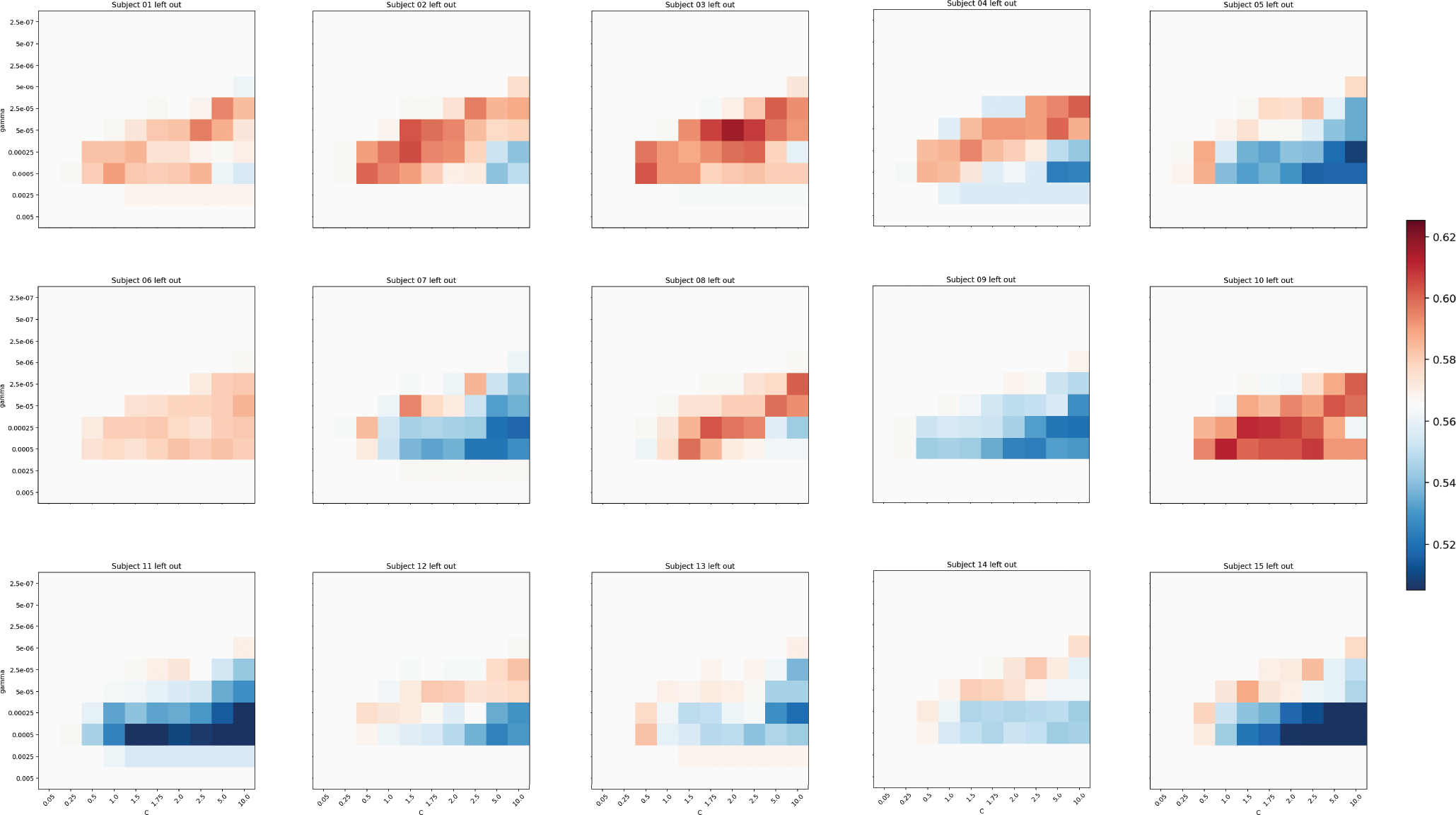
Cross-validation with a single held out subject to estimate parameters for the upper level performance single-trial SVM classifier. *c* values are displayed on the x-axis, and consisted of values: [0.05, 0.25, 0.5, 1, 1.5, 1.75, 2, 2.5, 5, 10]. *γ* values are displayed on the y-axis, and consisted of values: [2.5 × 10^−7^, 5 × 10^−7^, 2.5 × 10^−6^, 5 × 10^−6^, 2.5 × 10^−5^, 5 × 10^−5^, 2.5 × 10^−4^, 5 × 10^−4^, 2.5 × 10^−3^, 5 × 10^−3^].

**Fig. S9.**
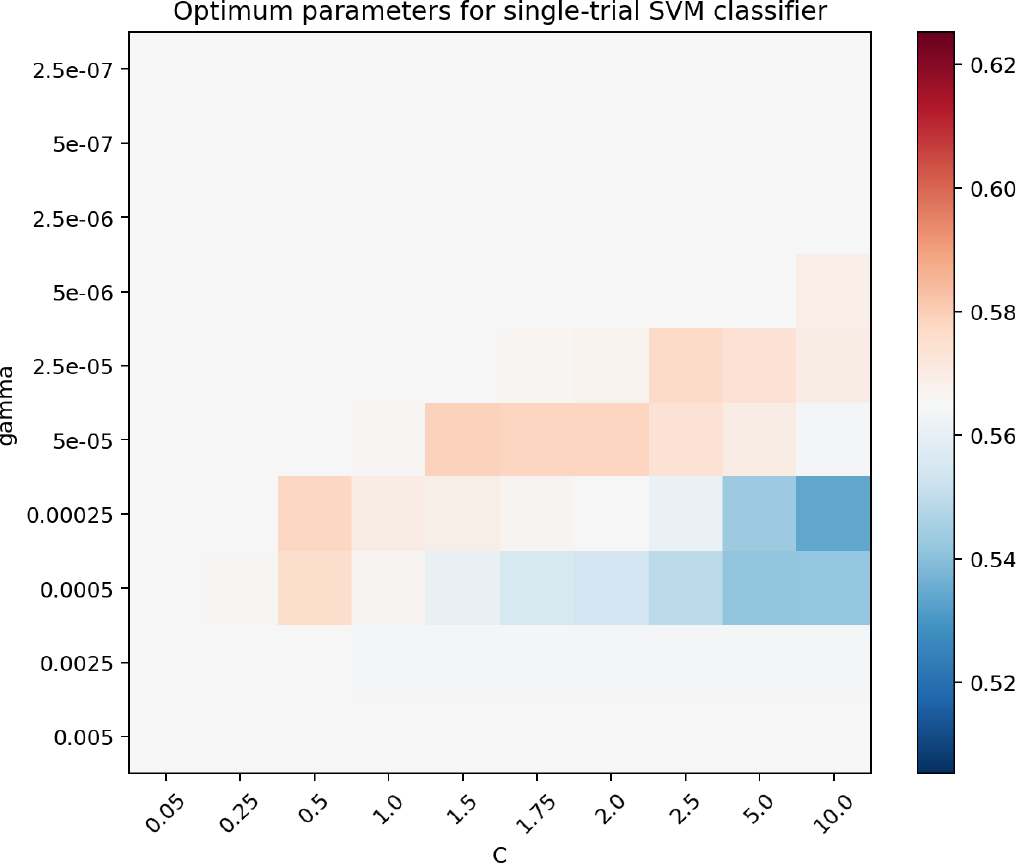
Upper level performance parameters for the single-trial SVM classifier based on the mean parameters for held out subjects in Figure S8.

**Fig. S10.**
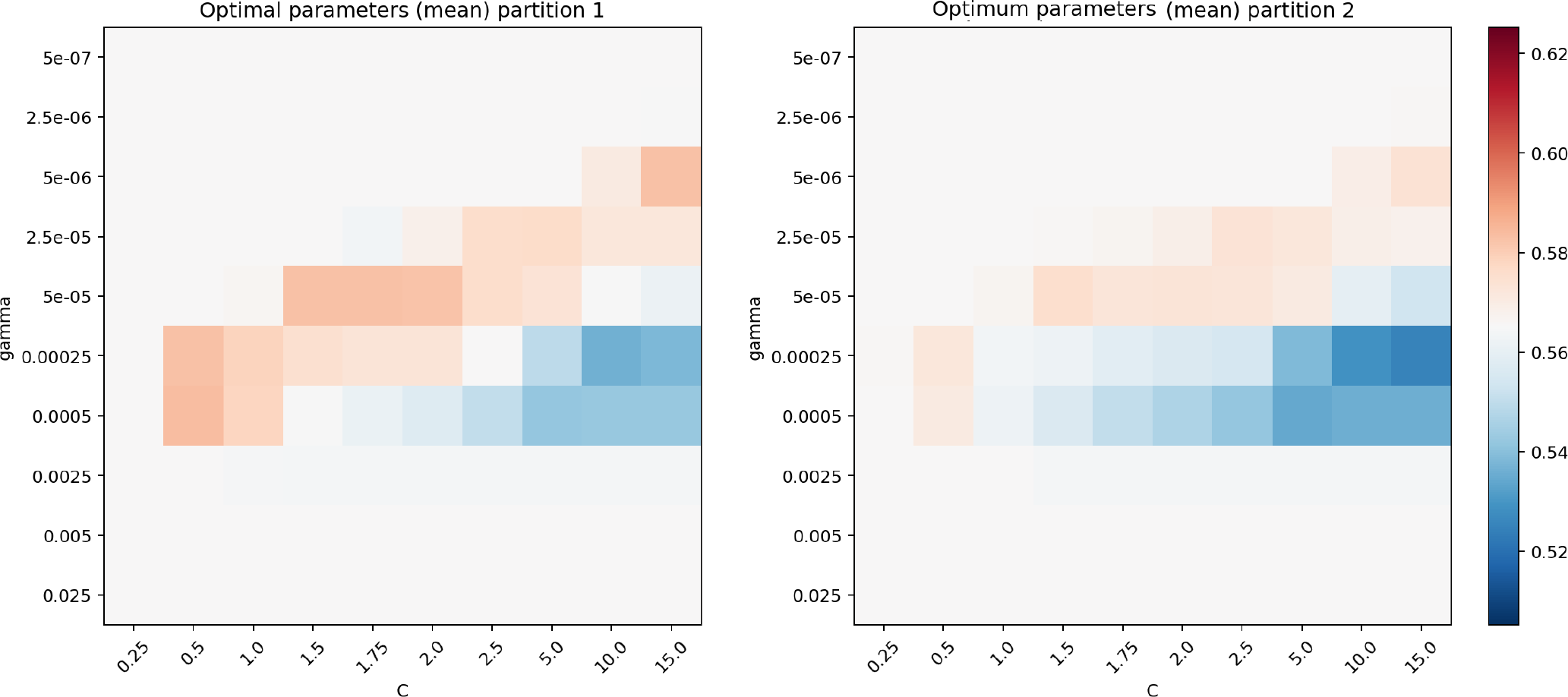
Optimum parameters for the single-trial SVM classifier based on the mean parameters of validation partition 1 (subjects 1-7) and partition 2 (subjects 8-15). Same cross-validation parameter values as in Figure S5.

